# Introducing THOR, a model microbiome for genetic dissection of community behavior

**DOI:** 10.1101/499715

**Authors:** Gabriel L. Lozano, Juan I. Bravo, Manuel F. Garavito Diago, Hyun Bong Park, Amanda Hurley, S. Brook Peterson, Eric V. Stabb, Jason M. Crawford, Nichole A. Broderick, Jo Handelsman

## Abstract

The quest to manipulate microbiomes has intensified, but many microbial communities have proven recalcitrant to sustained change. Developing model communities amenable to genetic dissection will underpin successful strategies for shaping microbiomes by advancing understanding of community interactions. We developed a model community with representatives from three dominant rhizosphere taxa: the Firmicutes, Proteobacteria, and Bacteroidetes. We chose *Bacillus cereus* as a model rhizosphere Firmicute and characterized twenty other candidates, including “hitchhikers” that co-isolated with *B. cereus* from the rhizosphere. Pairwise analysis produced a hierarchical interstrain-competition network. We chose two hitchhikers — *Pseudomonas koreensis* from the top tier of the competition network and *Flavobacterium johnsoniae* from the bottom of the network to represent the Proteobacteria and Bacteroidetes, respectively. The model community has several emergent properties—induction of dendritic expansion of *B. cereus* colonies by either of the other members and production of more robust biofilms by the three members together than individually. Moreover, *P. koreensis* produces a novel family of alkaloid antibiotics that inhibit growth of *F. johnsoniae,* and production is inhibited by *B. cereus*. We designate this community THOR, because the members are ***t***he ***h***itchhikers ***o***f the ***r***hizosphere. The genetic, genomic, and biochemical tools available for dissection of THOR provide the means to achieve a new level of understanding of microbial community behavior.

**IMPORTANCE:** The manipulation and engineering of microbiomes could lead to improved human health, environmental sustainability, and agricultural productivity. However, microbiomes have proven difficult to alter in predictable ways and their emergent properties are poorly understood. The history of biology has demonstrated the power of model systems to understand complex problems such as gene expression or development. Therefore, a defined and genetically tractable model community would be useful to dissect microbiome assembly, maintenance, and processes. We have developed a tractable model rhizosphere microbiome, designated THOR, containing *Pseudomonas koreensis, Flavobacterium johnsoniae,* and *Bacillus cereus,* which represent three dominant phyla in the rhizosphere, as well as in soil and the mammalian gut. The model community demonstrates emergent properties and the members are amenable to genetic dissection. We propose that THOR will be a useful model for investigations of community-level interactions.

## INTRODUCTION

Modern understanding of microbiomes has been accompanied by recognition of their vast complexity, which complicates their study and manipulation. Powerful –omics approaches that profile community features such as genomes, metabolites, and transcripts have illuminated the richness of many communities (1). These global portraits of complex communities have been complemented by genetic and biochemical dissection of much simpler communities and, in particular, binary interactions of one bacterial species with one host such as bacterial symbionts of legume roots (2) and squid light organs (3). Study of these systems has elucidated the pathways regulating interactions among bacteria and between bacteria and their environments.

The explosion of understanding of two-species interactions has generated a scientific thirst for more tools to attain mechanistic understanding of multi-species community behaviors. Model systems consisting of more than one microbial member include communities in flies (4, 5), the medicinal leech (6), and engineered systems (7). Key among the traits that demand more mechanistic studies are the components of community assembly and robustness, which is the ability to resist and recover from change (8, 9). Understanding these traits has particular value today as many researchers aim to modify microbial communities to achieve outcomes to improve human health, environmental sustainability, and agricultural productivity.

Over the last century, the challenge to alter microbial communities in predictable ways has stymied microbiologists. Examples include attempts to change the human gut microbiome by ingestion of yogurt or probiotics (10) or to alter plant microbiomes with inundative application of disease-suppressive microorganisms (11). Few treatments have induced long-term changes due to intrinsic community robustness.

Communities express emergent properties, which are traits that cannot be predicted from the individual members (12). For example, metabolic exchange between yeast and *Acetobacter* yields a mixture of volatile compounds attractive to *Drosophila* hosts that was not produced by either microbe in pure culture (13). Such higher-order or emergent properties of communities (14) might explain the difficulty in manipulating them. In addition, many communities contain functional redundancy that is likely to contribute to robustness.

Classical genetic analysis, defined as the isolation and study of mutants, in model organisms has advanced understanding of processes such as gene regulation in *E. coli* and the mouse, and development in flies and nematodes, but such reductionist genetic approaches are inaccessible in most microbial community-level analyses. A genetically tractable model system for studying microbial community assembly and robustness has the potential to transform our understanding of community processes. The scientific community recognizes the power of model systems in relation to microbial communities (15–17) and many groups have started to address this call (18–20). Importantly, common microbiome principles will only emerge with a diverse set of models to interrogate. Toward this end we developed a model system involving three bacterial species—*Pseudomonas koreensis, Flavobacterium johnsoniae,* and *Bacillus cereus*— which interact under field and laboratory conditions, are amenable to genetic analysis, and represent three major phyla in microbiomes on plant roots and in the human gut.

## RESULTS

### Source of community members

We sought a model community that is simple and contains genetically tractable species that likely interact under natural conditions, both competitively and cooperatively. We drew upon our previous work demonstrating the peculiar tendency of *B. cereus* to carry “hitchhikers” (21) when it is isolated from soybean roots. These biological hitchhikers are cryptic bacteria that become visible in culture only after apparently pure cultures of *B. cereus* are maintained at 4°C for several weeks (22). It is important to note that biological hitchhiking involves a physical association and is distinct from genetic hitchhiking which selects for neutral alleles (23). Most hitchhikers are members of the Bacteroidetes phylum with a small proportion from the Proteobacteria and Actinomycetes (22). We selected twenty-one candidates that include *B. cereus* UW85, twelve hitchhikers, and eight other isolates from the roots that harbored the hitchhikers (Fig. 1B-C). These isolates represented the four dominant phyla present in the rhizosphere: Firmicutes, Proteobacteria, Bacteroidetes, and Actinobacteria (24).

**FIG 1.**
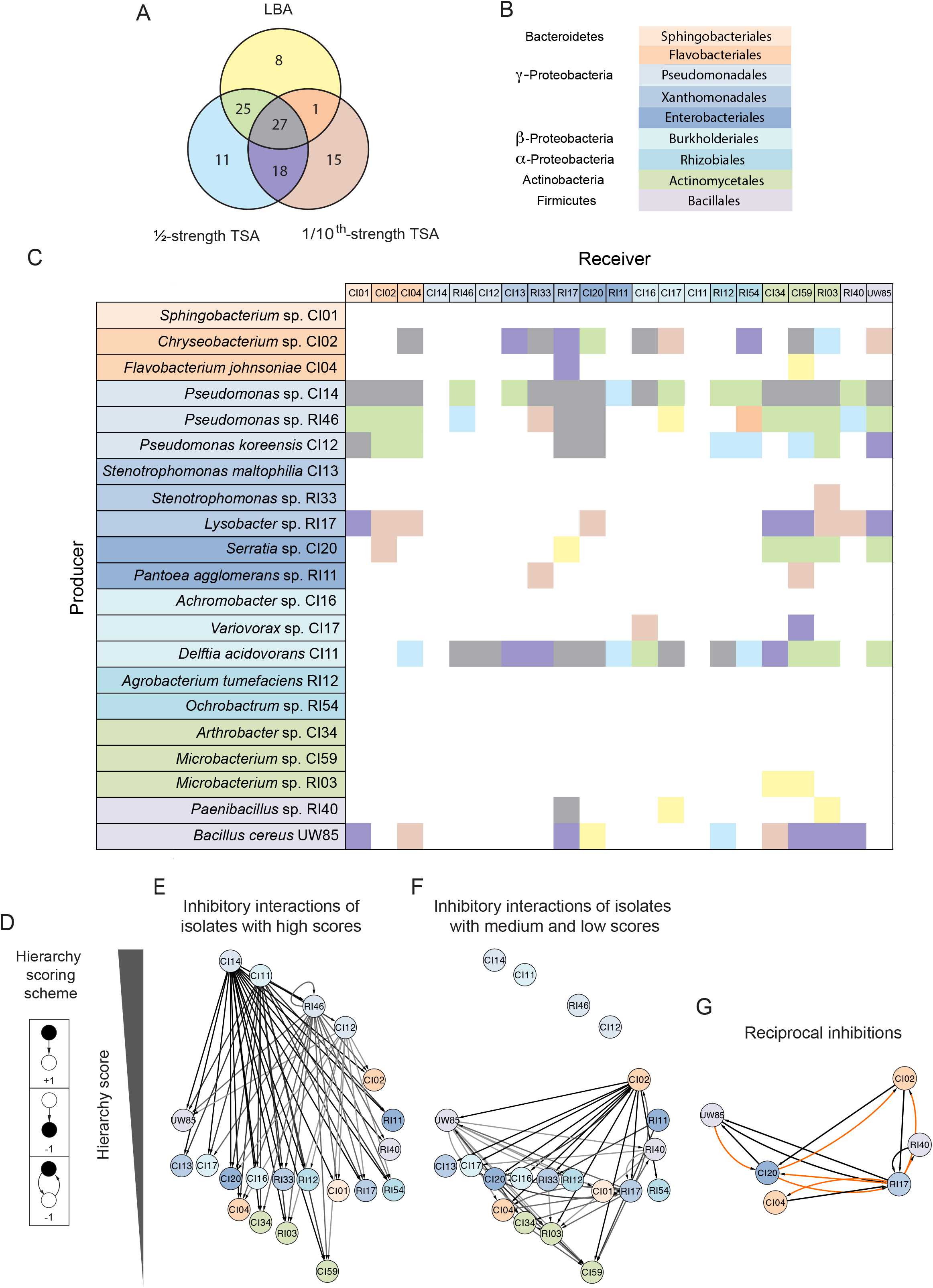
Network analysis of inhibitory interactions among rhizosphere isolates. (A) Venn diagram of the inhibitory interactions in isolates in three media: Luria-Bertani agar (LBA), ½-strength tryptic soy agar (TSA) and 1/10th-strength TSA. (B) Colors indicate phylogeny of isolates used in the inhibitory matrix and network. (C) Inhibitory interaction matrix between *B. cereus* UW85 and hitchhiker isolates in three media. Potential producers, which are isolates tested for the ability to inhibit others, are on the y-axis and receivers, which are isolates tested for inhibition by others, are on the x-axis. There are two different color codes used in (C). One indicates the phylogeny of isolates in the row and column title, and second one for the matrix results using the colors scheme shown in the Venn diagram corresponding to the medium in which the interaction appears. (D) Hierarchy scoring scheme used to organize the isolates in the hierarchy interaction network, in which black is the focal point. (E-F) Inhibitory interaction network organized by hierarchy score. (E) Inhibitory interactions generated from the isolates with high hierarchy scores. (F) Inhibitory interactions generated from the isolates with medium and low hierarchy scores. (G) Reciprocal inhibitory interactions observed in the inhibitory network. Orange indicates interactions observed in only 1 medium, and black indicates interactions observed in 2 or 3 media.

### High-order organization structures in an inhibitory interaction network of rhizosphere isolates

Competition plays an important role in microbial communities (25). To identify competitive interference interactions in our collection of twenty-one rhizosphere isolates, we evaluated them in pairs for inhibition of the other members in three different media. Of 105 inhibitory interactions detected, 71 (68%) were conserved in at least two growth conditions and 27 were conserved in all three conditions (Fig. 1A). From the 105 inhibitory interactions observed among the rhizosphere bacteria, we constructed a matrix (Fig. 1C). The strains display a high degree of interaction, indicated by high connectance (value of C = 0.24, which represents the fraction of all possible interactions or # interactions/# species^2^). On average, each isolate interacted with six other isolates, as either the source or target of inhibition. The interaction matrix appears to be producer determined with a negative sender-receiver asymmetry value of (Q) = -0.31 (26). This suggests that the structure of the inhibitory networks is more controlled by the producers and their secreted antibiotics than by the tester strains. Connectance values reported previously for diverse food-web structures are between 0.026 and 0. 315 (27), and the strains tested in this study had a value of 0.24, indicating high connectance.

To detect higher order community organization mediated by inhibitory interactions, we used a hierarchy-scoring scheme in which the ability to inhibit was given a positive score and sensitivity generated a negative score (Fig. 1D). We observed inhibitory hierarchical interactions in which the top isolates of the network inhibit isolates that receive a medium score, and in turn these medium-scoring isolates inhibit isolates that receive a lower score (Fig. 1E-F).

The four isolates that received the highest score, *Delftia acidovorans* and three *Pseudomonas* spp., were responsible for 58 (55%) of the inhibitory interactions (Fig. 1E). Non-hierarchical interactions were infrequent (6%), and these were largely reciprocal interactions between six isolates with middle and low scores, such as *Lysobacter* sp. RI17 (Fig. 1G), which is inhibited by five isolates under almost all conditions. Reciprocal inhibition by *Lysobacter* sp. against these strains appeared predominantly in the lower nutrient condition (1/10^th^-strength TSA).

### *P. koreensis* isolate inhibits members of the Bacteroidetes in root exudate

Three of the strains inhibited most of the other isolates, placing them at the top of the inhibitory network. These strains were all members of the *P. fluorescens* complex. They are the only members of the collection that consistently inhibit *Bacteroidetes* isolates, which is the most abundant phylum among the co-isolates of *B. cereus,* making the inhibitory activities of these *P. fluorescens* members of particular interest. In addition, members of the *P. fluorescens* group have been shown to alter the structure of microbial communities on roots, making the inhibition of Bacteroidetes of interest to understand communities (28). To determine whether the three *Pseudomonas* strains inhibit members of the *Bacteroidetes* similarly, we tested them against *F. johnsoniae* CI04. *Pseudomonas* sp. CI14 and *Pseudomonas* sp. RI46 inhibited *F. johnsoniae* CI04 growth in standard media and *P. koreensis* CI12 inhibited *F. johnsoniae* CI04 only in root exudate (Fig. 2A). Against a panel of diverse rhizosphere isolates grown in root exudate, *P. koreensis* CI12 inhibited primarily Bacteroidetes strains (Fig. 2B).

**FIG 2.**
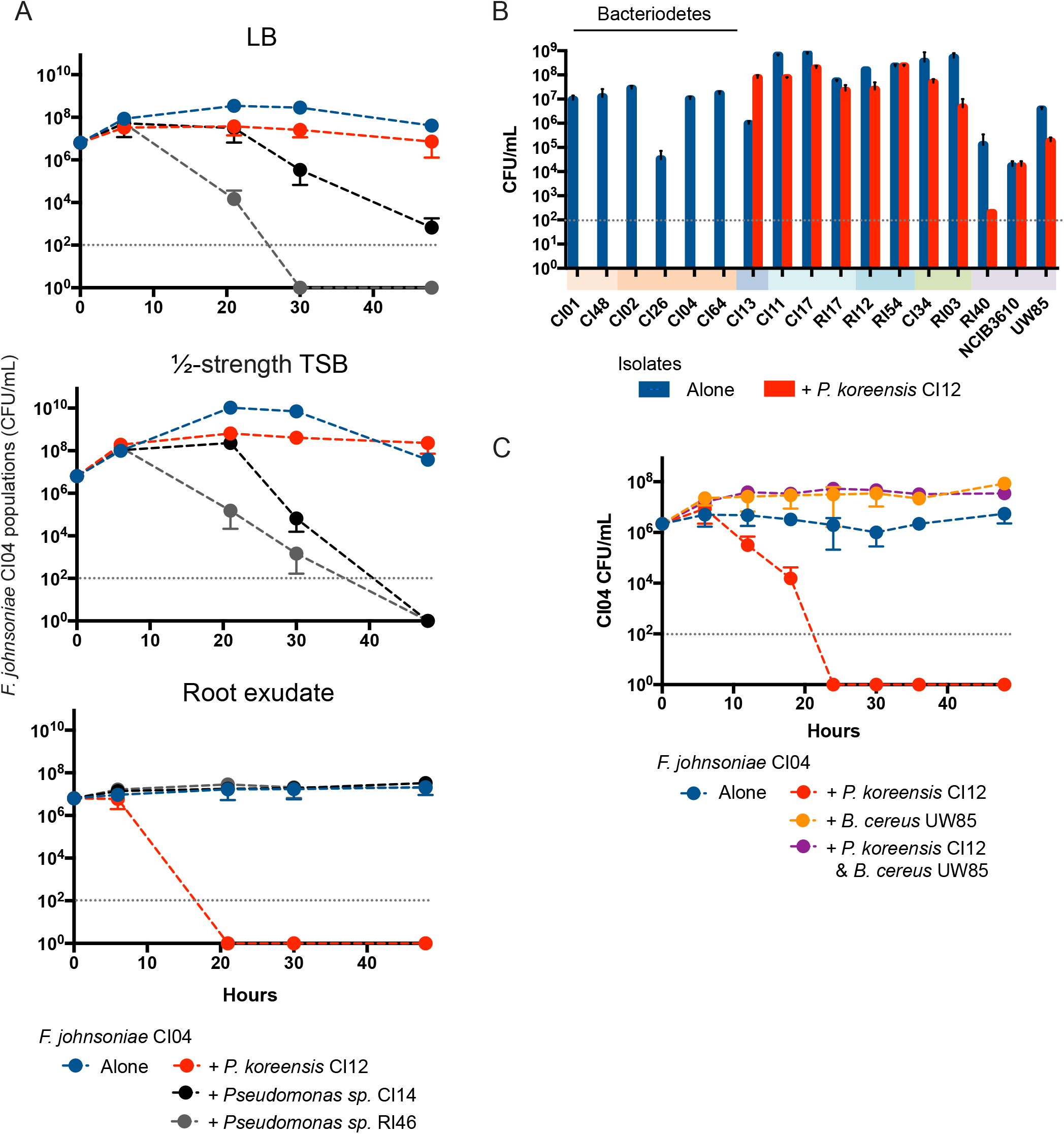
Co-culture of rhizosphere isolates with *Pseudomonas spp.* and *B. cereus* UW85. (A) Competition experiments between *F. johnsoniae* CI04 and either *Pseudomonas* sp. CI14, *Pseudomonas* sp. RI46, or *P. koreensis* CI12, in three media: Luria-Bertani broth (LB), ½-strength tryptic soy broth (TSB) and soybean root exudate. (B) Rhizosphere isolates were grown alone or in co-culture with *P. koreensis* CI12 in soybean root exudate. Colored bars under x-axis indicate phylogenetic groups as in Fig. 1. (C) *F. johnsoniae* CI04 grown alone, in co-culture with *P. koreensis* CI12 or *B. cereus* UW85, and in triple culture with *P. koreensis* CI12 and *B. cereus* UW85. Gray dotted line, limit of detection.

### *B. cereus* protects *F. johnsoniae* from *P. koreensis* by modulating levels of koreenceine metabolites

To explore the ecology of *B. cereus* and its hitchhikers, *P. koreensis* CI12 and *F. johnsoniae* CI04, we added *B. cereus* UW85 to the co-culture of the hitchhikers. Unpredictably, *B. cereus* UW85 enabled growth of *F. johnsoniae* CI04 in co-culture with *P. koreensis* CI12 (Fig. 2C) without affecting growth of *P. koreensis* CI12 (Fig. S1). *F. johnsoniae* protection is dependent upon *B. cereus* arrival at stationary phase (Fig. S2). *B. subtilis* NCIB3160, a well-studied spore-forming bacterium, also protected *F. johnsoniae* in a stationary phase-dependent manner (Fig. S2). Among three *B. subtilis* NCIB3160 mutants affected in transcriptional regulators related to stationary phase (29), only the mutant affected in *spo0H,* which controls the early transition from exponential to stationary phase, was unable to protect *F. johnsoniae* from *P. koreensis* (Fig. 3). A *spo0H* mutant in *B. cereus* UW85 also did not protect *F. johnsoniae* from *P. koreensis* (Fig. 3).

**FIG 3.**
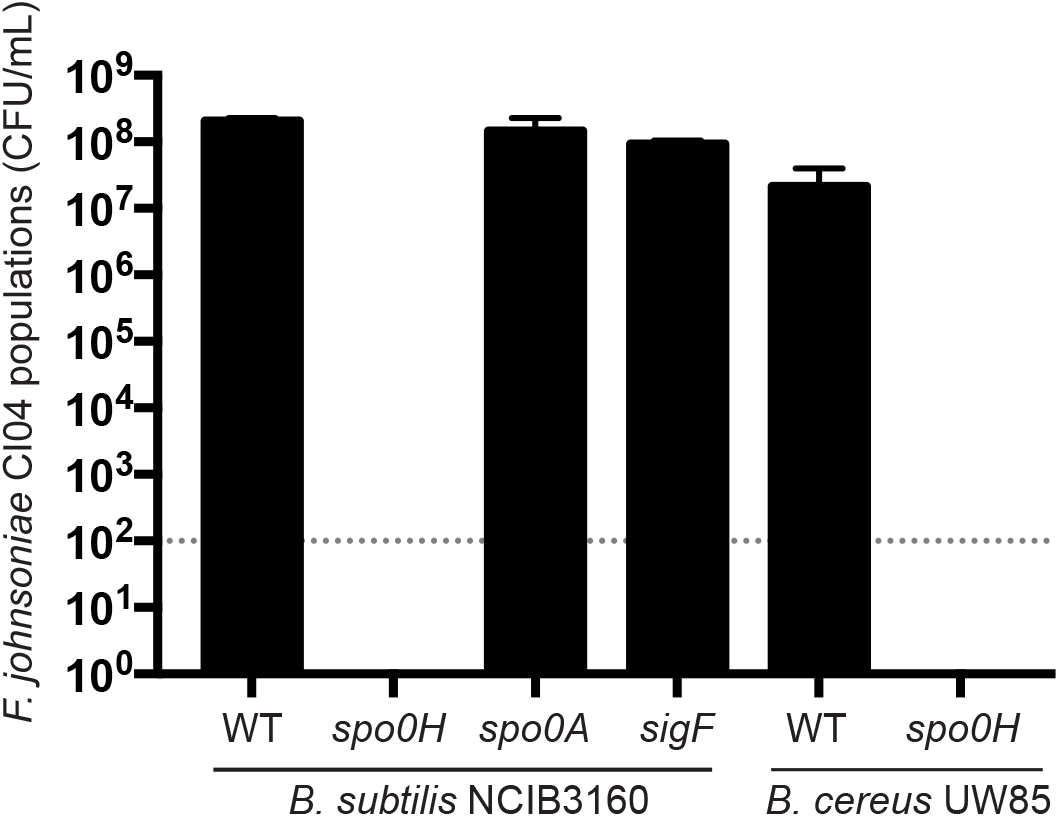
Effect of *Bacillus* spp. on populations of *F. johnsoniae* in the presence of *P. koreensis.* Triple culture of *P. koreensis* CI12, *F. johnsoniae* CI04, and either *B. subtilis* NCIB3160 or *B. cereus* UW85, or their mutants. Gray dotted line, limit of detection.

We recently described a *P. koreensis* family of bacterial alkaloids—koreenceine A, B, and C—that influence growth of *F. johnsoniae* in root exudates (30). Cell-free filtrates of co-cultures of *P. koreensis* and *F. johnsoniae* inhibit *F. johnsoniae* growth, whereas filtrates of *P. koreensis* cultured alone or with both *F. johnsoniae* and *B. cereus* do not inhibit *F. johnsoniae* growth (Table 1). The levels of koreenceine A, B, and C were higher when *P. koreensis* was co-cultured with *F. johnsoniae* than when it was cultured alone (Table 1). Addition of *B. cereus* to co-cultures of *P. koreensis* and *F. johnsoniae* severely reduced accumulation of koreenceine A and C with a minor effect on the level of koreenceine B. The *B. cereus* Δ*spo0H* mutant did not protect *F. johnsoniae* in triple culture, nor did it reduce accumulation of koreenceine A and C as substantially as did wild type *B. cereus* (Table 1). We propose that *B. cereus* protects *F. johnsoniae* by selectively reducing the levels of koreenceine A and C accumulated by *P. koreensis*.

**TABLE 1.**
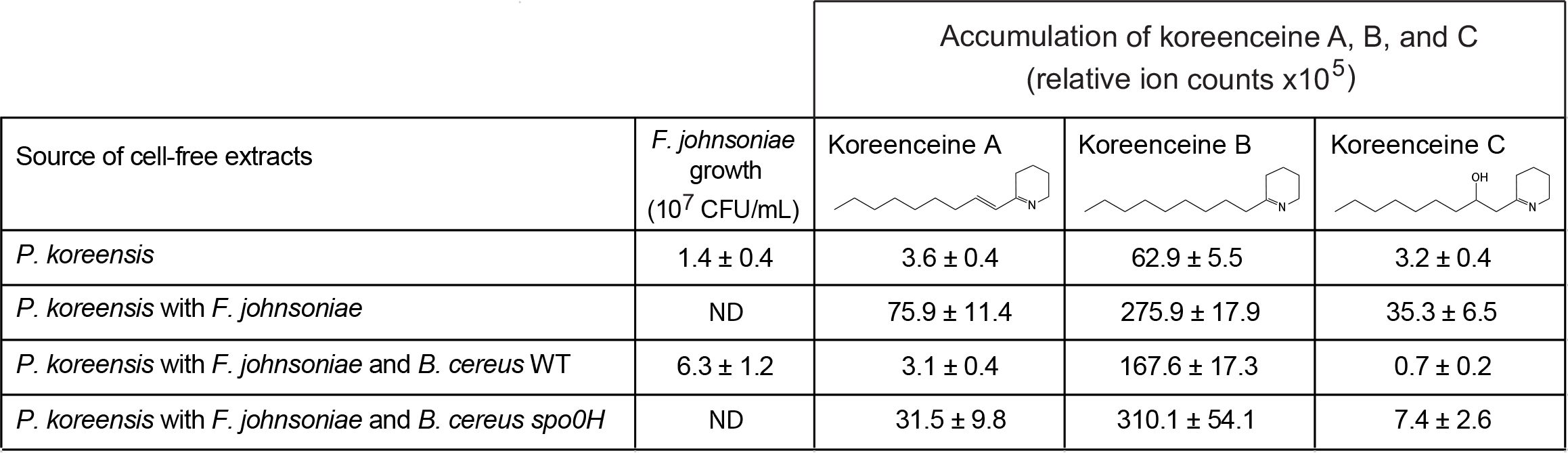
Accumulation of koreenceine A, B, and C in co-culture with *F. johnsoniae* or *B. cereus.* Koreenceine A, B, and C concentrations are expressed as ion counts from LC/HR-ESI-QTOF-MS analysis from cell-free filtrates of cultures of *P. koreensis* CI12 grown alone, with *F. johnsoniae* CI04, *F. johnsoniae* CI04 and *B. cereus* UW85 wild type, or with *F. johnsoniae* CI04 and *B. cereus* UW85 *spo0H. F. johnsoniae* CI04 population growth in the corresponding conditions in CFU/mL. ND, not detected.

|  |  | Accumulation of koreenceine A, B, and C<br>(relative ion counts x10<sup>5</sup>) |  |  |
| --- | --- | --- | --- | --- |
| Source of cell-free extracts | *F. johnsoniae*<br>growth<br>(10<sup>7</sup> CFU/mL) | Koreenceine A | Koreenceine B | Koreenceine C |
| *P. koreensis* | 1.4 ± 0.4 | 3.6 ± 0.4 | 62.9 ± 5.5 | 3.2 ± 0.4 |
| *P. koreensis* with *F. johnsoniae* | ND | 75.9 ± 11.4 | 275.9 ± 17.9 | 35.3 ± 6.5 |
| *P. koreensis* with *F. johnsoniae* and *B. cereus* WT | 6.3 ± 1.2 | 3.1 ± 0.4 | 167.6 ± 17.3 | 0.7 ± 0.2 |
| *P. koreensis* with *F. johnsoniae* and *B. cereus spo0H* | ND | 31.5 ± 9.8 | 310.1 ± 54.1 | 7.4 ± 2.6 |

### Rhizosphere isolates modulate *B. cereus* colony expansion

Among twenty rhizosphere isolates, six induced *B. cereus* patches to expand in a dendritic pattern when plated on a lawn of the corresponding isolate on 1/10^th^-strength TSA (Fig. 4). *B. cereus* was the only isolate of the collection that displayed colony expansion. A similar motility pattern has been observed in *B. cereus* translocating in an artificial soil microcosm (31), suggesting that this motility might be important in adapting to the soil environment. In pure culture, *B. cereus* colonies expanded in an irregular pattern with dense branches and asymmetrical bulges, whereas on lawns of the six isolates, including *P. koreensis* and *F. johnsoniae,* expansion was greater and radially symmetrical (Fig. 4, Fig. 5A, Table S1). Colonies of *B. cereus* spread into the neighboring colonies of *F. johnsoniae* CI04 and *P. koreensis* CI12 when the two strains were grown in close proximity on plates (Fig. 5B). *B. cereus* spread both around and across *P. koreensis* CI12, and across, but not around, *F. johnsoniae* CI04. *B. cereus* did not display colony expansion when in contact with or proximal to *Paenibacillus* sp. RI40 (Fig. 5).

**FIG 4.**
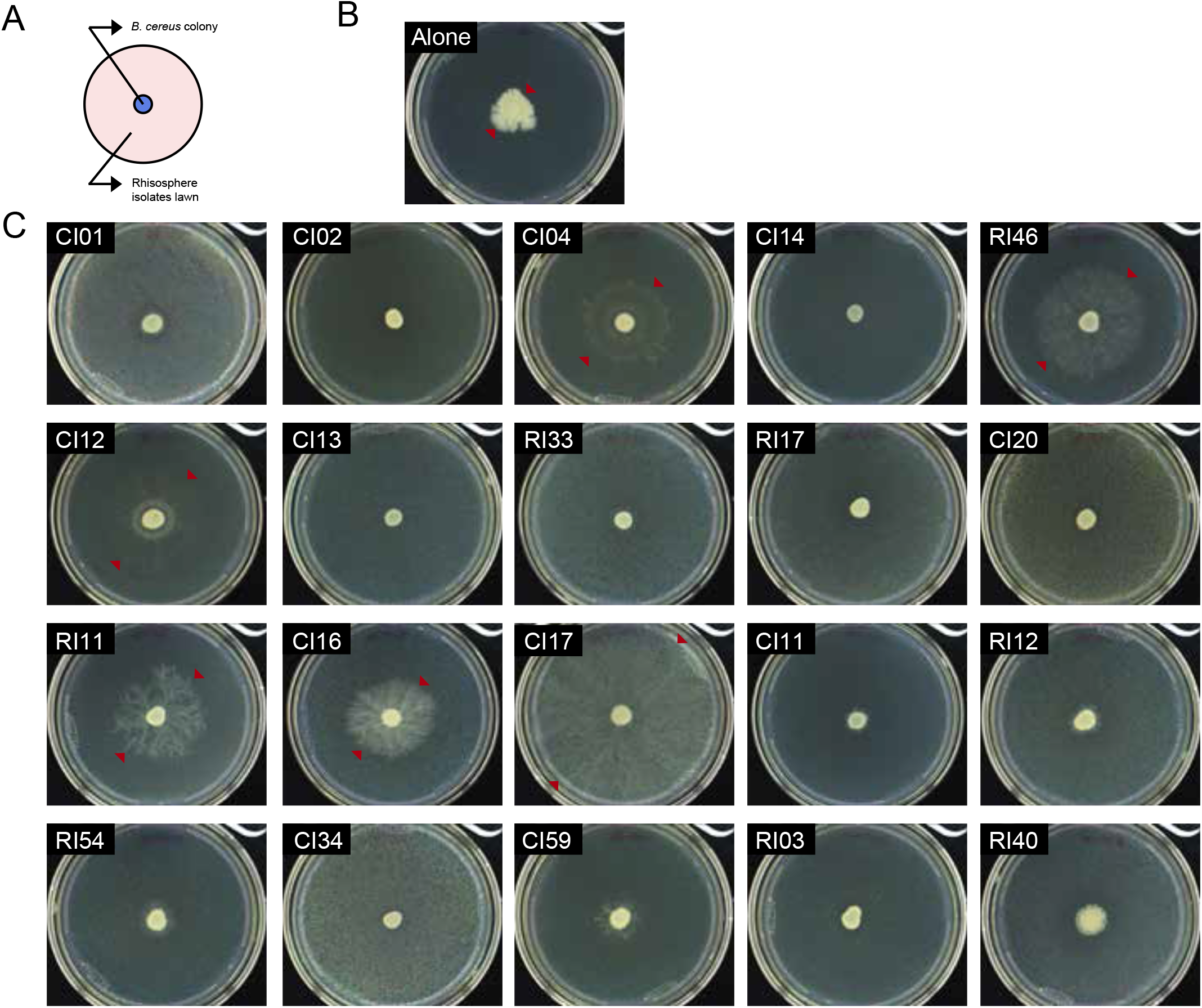
*B. cereus* UW85 colony expansion in the presence of the rhizosphere isolates. (A) Schematic representation of the spread-patch plates. (B) *B. cereus* UW85 grown alone. (C) *B. cereus* UW85 grown on a lawn of each member of the community. Photographs taken after 4 days at 28°C. Arrows indicate the limits of the *B. cereus* colony.

**FIG 5.**
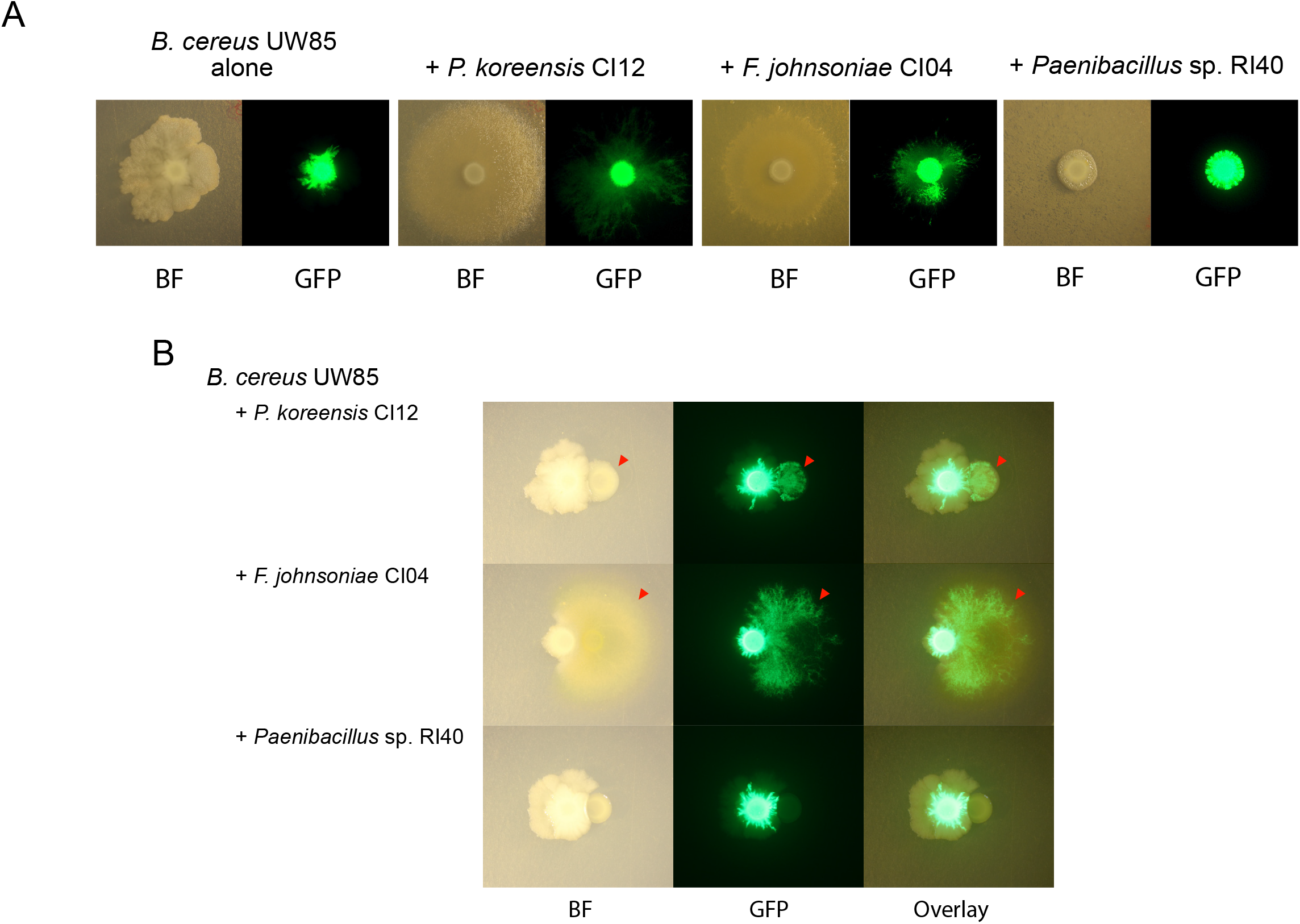
Effect of community members on *B. cereus* UW85 colony expansion. (A) *B. cereus* UW85 plasmid-dependent GFP strain grown alone or on a lawn of *P. koreensis* CI12, *F. johnsoniae* CI04, or *Paenibacillus* sp. RI40. Bright-field (BF) and GFP imaging of colonies five days after inoculation. (B) *B. cereus* UW85 GFP strain grown in close proximity to a colony of *P. koreensis* CI12, *F. johnsoniae* CI04, or *Paenibacillus* sp. RI40. Arrows indicate *B. cereus* UW85 expansion over colonies of the other isolates. Bright-field, GFP channel, and overlay of the two channels of plates after 2 days growth (*F. johnsoniae* CI04) and after 5 days growth (*P. koreensis* CI12 and *Paenibacillus* sp. RI40).

Rhizosphere isolates modulate *Pseudomonas* biofilm formation. Among the hitchhikers and rhizosphere isolates, *Pseudomonads spp.* isolates produced the most robust biofilms (Fig. 6A). In pairwise tests, poor biofilm producers changed the behavior of isolates of *Pseudomonas spp.* (Fig. 6B-D). When alone, *P. koreensis* CI12 produced maximum biofilm at 18 h, after which the biofilm began to dissociate (Fig. S3). *F. johnsoniae* CI04 and *B. cereus* UW85 each increased the maximum biofilm formed by *P. koreensis* CI12 at 18 h, and reduced the rate of biofilm dissociation (Figure 6E-F). A mixture of all three strains followed the same pattern as the pairs, although the triple mixture maintained more biofilm at 36 hours (p<0.01) (Fig. 6F), suggesting that *F. johnsoniae* CI04 and *B. cereus* UW85 together sustain the *P. koreensis* CI12 biofilm longer than either alone.

**FIG 6.**
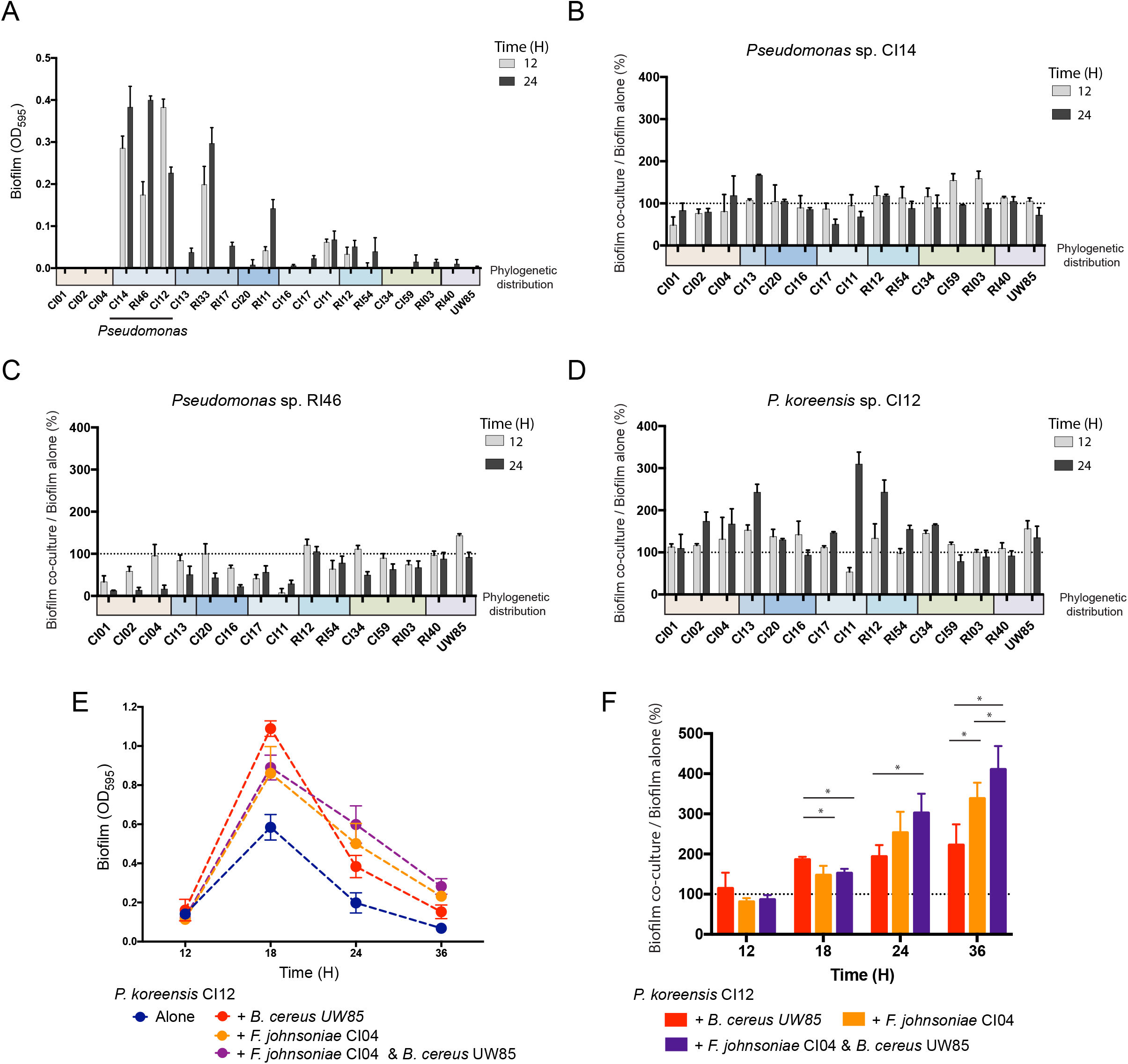
Biofilm formation by rhizosphere isolates. Biofilm was quantified by measuring the OD595 after staining with crystal violet. (A) Crystal violet quantification of biofilm formation for each of the 21 isolates at 12 and 24 hours after inoculation. (B-D) *Pseudomonas* biofilm production when grown with a poor biofilm producer normalized against *Pseudomonas* biofilm in pure culture at 12 and 24 hours (B) *Pseudomonas* sp. RI46 (C) *Pseudomonas* sp. CI14 (D) *P. koreensis* CI12. (E) Biofilm formation by *P. koreensis* CI12 growing alone, in co-culture with either *F. johnsoniae* CI04 or *B. cereus* UW85, and in triple culture at 12, 18, 24, and 36 hours. (F) *P. koreensis* CI12 biofilm production when grown with two other isolates normalized against *P. koreensis* growth in pure culture. * indicates p<0.01. Colored bars under x-axis indicate phylogenetic groups as in Fig. 1. Gray dotted line, limit of detection.

## DISCUSSION

We constructed a model community built upon microbes isolated from the soybean rhizosphere. We selected *B. cereus* as the first member of the community because of its ubiquitous distribution in soil and on roots (32) and its influence on the rhizosphere (33–36). Approximately 3 to 5% of *B. cereus* colonies isolated from roots carry “hitchhikers”—other species that are only visible in culture over time in cold storage (22). *B. cereus* and its hitchhikers are derived from the same habitat and appear to have a physically intimate association, suggesting that their interactions in culture are relevant to the natural community; therefore, the second and third candidates for the model community were chosen from among the hitchhikers.

The second candidate was *F. johnsoniae,* a member of the Bacteroidetes, the most abundant group of hitchhikers. *B. cereus* enables *F. johnsoniae* to grow in soybean and alfalfa root exudate by providing fragments of peptidoglycan as a carbon source (22). The third candidate for the model community was *P. koreensis,* which inhibits growth of *F. johnsoniae* in soybean root exudates, but not when *B. cereus* is present. Other interactions among these three species include modulation of *B. cereus* colony expansion by *F. johnsoniae* and *P. koreensis* and enhancement of *P. koreensis* biofilm formation by *B. cereus* and *F. johnsoniae.* These three candidates are genetically tractable, their genomes have been sequenced (21, 37, 38), they represent three phyla that dominate the rhizosphere and other host-associated communities (39-44), and they display both competitive and cooperative interactions. We designate the model community containing *B. cereus, F. johnsoniae,* and *P. koreensis* as “THOR,” to indicate the members are *the* hitchhikers of the rhizosphere.

THOR has two emergent properties, colony expansion and biofilm formation, that are increased by the complete community and could not be predicted from the behavior of the individuals. Each are observed with several combinations of community members, demonstrating functional redundancy in the system. *B. cereus* colony expansion likely reflects other bacteria affecting *B. cereus* motility. To our knowledge, this is the first report to indicate *B. cereus* motility influenced by social interactions. Motility influenced by social interactions is a rapidly growing field of research that has already demonstrated diverse mechanisms by which bacteria modulate motility in other species, including diffusible metabolites and cell-to-cell contact (45, 46). The *B. cereus* dendritic growth patterns described here have been associated with sliding motility in semi-solid agar under low-nutrient conditions (47), and in artificial soil microcosms (31). Sliding is a passive appendage-independent translocation mechanism mediated by expansive forces of a growing colony accelerated by biosurfactants (48). Future experiments will determine whether *B. cereus* colony expansion over a bacterial lawn is mediated by sliding and whether the inducing bacteria enable this motility with biosurfactants.

Biofilm produced by *P. koreensis* CI12 was augmented and sustained by other THOR members. *P. koreensis* CI12 is a member of the *P. fluorescens* complex, which contains many members that promote growth and suppress disease of plants, processes often dependent upon biofilm formation (49). The community modulation of *P. koreensis* CI12 biofilm formation and persistence could be a strategy to maintain bacteria in the rhizosphere. The social interactions among the members of the THOR model community provides an experimental system to probe mechanisms of *P. koreensis* biofilm assembly and disassembly and its role in *P. koreensis* lifestyle within a rhizosphere community.

In addition to having emergent community properties, THOR members interact in several ways that are common in communities. *B. cereus* increases *F. johnsoniae* growth through nutritional enhancement (22) and protects it from growth inhibition by *P. koreensis,* illustrating how pairwise interactions can be modulated by other members of the community, a phenomenon observed previously in synthetic communities (50, 51). Growth interference and enhancement have been considered rare in naturally occurring communities because these behaviors often cannot be predicted from pairwise interactions (52). Our results further reinforce the importance of community modulation of behaviors observed in pairwise studies.

THOR was constructed from a collection of 21 rhizosphere isolates that share several properties such as high interactivity and high-order organization mediated by antagonistic interactions with previously proposed model communities (26, 53, 54). We show that these properties can originate from phylogenetically diverse bacteria. We propose that the high microbial diversity detected in soil and rhizosphere communities could be in part achieved by hierarchical inhibition coupled with modulation of inhibition.

A genetically tractable community with defined composition in a controlled environment offers the opportunity to dissect the mechanisms by which communities are established, function, and maintain their integrity in the face of perturbation. Rigorous testing of the numerous mechanistic hypotheses about community behavior that have been generated by -omics analyses requires systems in which variables can be isolated. This is offered in genetically tractable systems in which the functions of individual genes can be established through mutant analysis, and the impact of environmental factors can be studied by manipulating each variable. Therefore, model systems that can be fully dissected need to be part of the arsenal of tools to advance microbial ecology to a new platform of experimental power and causal inference.

We present THOR as a simple, multi-phylum, genetically tractable system with diverse community characteristics, some of which are the result of emergent properties. To capture the impact of multi-organism interactions and emergent properties, communities need to be studied as single genetic entities. Metagenomics introduced the concept of the community as the unit of study for genomes; similarly, “metagenetic” analysis will apply genetic analysis at the community level for mechanistic understanding (55). Such understanding will be key to designing interventions to achieve outcomes in the health of humans, the environment, and the agroecosystem.

## ACKNOWLEDGMENTS

We gratefully acknowledge Jennifer Heinritz for her assistance with microscopy. This research was supported by the Office of the Provost at Yale University, by funding from the Wisconsin Alumni Research Foundation through the University of Wisconsin–Madison Office of the Vice Chancellor for Research and Graduate Education, and by NSF grant MCB-1243671.

## MATERIALS AND METHODS

### Bacterial strains and culture conditions

*B. cereus* UW85 and 20 co-isolates and rhizosphere isolates were reported previously (22)(Table S2). *B. subtilis* NCIB3160 WT, *spo0A, spo0H* and *sigF* were a gift from Roberto Kolter at Harvard University. Bacterial strains were propagated on 1/10^th^-strength tryptic soy agar (TSA) and grown in liquid culture in ½-strength tryptic soy broth (TSB) at 28°C with vigorous shaking. *Bacillus* spores were quantified by plating on 1/10^th^ TSA after heating at 80°C for 10 min.

### Production of root exudates

Soybean seeds were surface disinfected with 6% sodium hypochlorite for 10 min, washed with sterilized deionized water, transferred to water agar plates, and allowed to germinate for three days in the dark at 25°C. Seedlings were grown in a hydroponic system using modified Hoagland's plant growth solution (56). Root exudate was collected after 10 days of plant growth in a chamber (12-h photoperiod, 25°C), filter sterilized and stored at -20°C. An amino acid mix of equal parts alanine, aspartate, leucine, serine, threonine, and valine was added to the root exudate at a final concentration of 6 mM.

### Generating an inhibitory interaction network between rhizosphere bacteria

The presence or absence of inhibitory interactions between strains in our collection was evaluated following a modified spread-patch method. Strains were grown individually for 20 h. One-mL aliquots of cultures of each strain were centrifuged (6000 × g, 6 min), resuspended in 1 ml of the same medium (undiluted cultures), and a 1:100 dilution of each strain was prepared in the same medium (diluted culture). Inhibitory interactions were evaluated in three different media plates: Luria-Bertani agar, 1/2-strength TSA, and 1/10^th^-strength TSA. Plates were spread with 100 μL of the diluted cultures and spotted with 10 μL of the undiluted cultures, with four strains per plate. Plates were then incubated at 28°C and inspected for zones of inhibition after two days. A network of inhibitory interactions was then generated using the inhibitory interaction matrix that summarize the detected interactions in the three conditions evaluated, where each node of the network represents one of the bacterial strains and each edge represents growth inhibition of the target. A simple hierarchy scoring was created to assign hierarchy levels based on Wright & Vetsigian, 2016 (54). Each strain was assigned one “reward” point for inhibiting another strain and one “penalty” point for each strain that inhibited it. One penalty point was also assigned for reciprocal interactions. Networks were visualized using Cytoscape software (57). The sender-receiver asymmetry (Q) was calculated from the inhibitory interaction matrix as reported by Vetsigian et al., 2011 (26).

### Competition assays in liquid culture

Strains were grown individually for 16 to 20 h. A one-mL sample was removed from each overnight culture and the cells were washed once and resuspended in phosphate-buffered saline (PBS). Culture medium was inoculated with ~10^6^ *F. johnsoniae* cells mL^-1^, and ~10^7^ *Pseudomonas* cells mL^-1^. *B. cereus* and *B. subtilis* were inoculated at densities between ~10^4^ cells mL^-1^ to ~10^7^ cells mL^-1^, depending upon the experiment. When evaluating the susceptibility of each strain to *P. koreensis* CI12 inhibition in root exudate, all strains except CI12 were inoculated at a final density of 1/1000 their overnight culture density. Cultures were incubated with agitation for two days at 28°C. Samples were withdrawn periodically to evaluate bacterial growth by serial dilution and plating. The initial densities were determined on either LB or LB containing kanamycin (10 μg mL^-1^). At every other time point, *Pseudomonas* sp. colonies were counted on LB plates, *Bacillus* sp. were selected on LB plates containing polymyxin B (4 μg mL^-1^), and all other strains were selected on LB plates containing kanamycin (10 μg mL^-1^). Plates were incubated at 28°C for two days.

### Chromosomal deletion of *spo0H* in *B. cereus*

The gene encoding the Spo0H sigma factor was deleted using a chromosomal integration vector with a thermosensitive origin of replication that introduces the deletion with no marker. Construction of the *spo0H* deletion cassette was accomplished by a modified version of overlap extension (OE) PCR strategy. Fragments one kb upstream and one kb downstream of the *spo0H* gene were amplified using primers mut_spo0HA1/mut_spo0HA2 and mut_spo0HB1/mut_spo0HB2 respectively (Table S3). The PCR products were cloned in pENTR/D-TOPO, generating pmut_spo0H_ENTR. Primers mut_spo0HA1 and mut_spo0HB2 were designed to include a BamHI site in the 5’ region to allow transfer. The *spo0H* deletion construct was recovered from pmut_spo0H_ENTR using BamHI, and cloned in the BamHI site of pMAD, generating pmut_spo0H_MAD in *E. coli* GM2929. Gene replacement was carried out in a manner similar to the method described previously (58). Briefly, pmut_spo0H_MAD was introduced into *B. cereus* UW85 by electroporation in 0.2-cm cuvettes with a Gene Pulser (BioRad Laboratories) set at 1.2 kV. The surviving cells were cultured on 1/2-strength TSA plates with erythromycin (3 μg mL^-1^) and X-Gal (50 μg mL^-1^) at 28°C. Strains that had undergone the single recombination event were selected on plates containing erythromycin (3 μg mL^-1^) at 40.5°C. To select for a second crossover event, single recombinant clones were grown at 28°C in nonselective media, diluted into fresh medium and then grown at 40.5°C, and plated for single colonies on 1/2-strength TSA with X-Gal (50 μg mL^-1^) at 40.5°C. White colonies, which were putative double recombinants, were confirmed by PCR using mut_spo0HA1 and mut_spo0HB2 for the deletion of *spo0H.*

### Identification of koreenceine metabolites produced by *P. koreensis* CI12 in the presence of other rhizosphere members

*P. koreensis* CI12 was grown alone, in co-culture or in triple culture with other rhizosphere members in root exudate at 28°C with agitation. Five mL of cultures were centrifuged (6000 × *g*, 6 min), and supernatants were filtered using a 0.22-μm pore size polyethersulfone (PES) membrane filter (Millipore). The cell-free cultures were mixed with 6 mL of 2-butanol. Six mL of the organic phase was concentrated using a GeneVac EZ-2 plus (SP Scientific). Crude extracts were resuspended in methanol and analyzed on an Agilent 6120 single quadrupole liquid chromatography-mass spectrometry (LC/MS) system (Column: Phenomenex kinetex C_18_ column, 250 × 4.6 mm, 5 μm; flow rate: 0.7 mL min^-1^ mobile phase composition: H_2_O and acetonitrile (ACN) containing 0.1% trifluoroacetic acid (TFA); method: 0-30 min, 10-100% ACN; hold for 5 min, 100% ACN; 1 min, 100-10% ACN). High-resolution electrospray ionization mass spectrometry (HR-ESIMS) data were obtained using an Agilent iFunnel 6550 Q-TOF (quadrupole-time-of-flight) mass spectrometer fitted with an electrospray ionization (ESI) source coupled to an Agilent (USA) 1290 Infinity high performance liquid chromatography (HPLC) system.

### *B. cereus* motility assay

*B. cereus* motility was evaluated using the same modified spread-patch method described above. We used *B. cereus* UW85/pAD123_31-26 as a GFP reporter (59). The images were captured using a custom Macroscope. The detector was a Canon EOS 600D Rebel T3i equipped with a Canon EFS 60mm Macro lens. GFP was excited with an LED using a 470/40 filter and collected with a 480/30 filter. Remote control of the camera and LED was achieved using custom software.

### Microtiter plate biofilm assays

The ability of the rhizosphere community to form a biofilm was estimated in 96-well polystyrene microtiter plates as described previously (60) with some modification. Briefly, strains were grown at 28°C for 20 h; cultures were centrifuged (6000 × g, 6 min) and resuspended in sterile 10 mM NaCl to an optical density (OD) of 0.004 for *Pseudomonas* spp. and 0.001 for the other isolates. Cell suspensions were placed in sterile flat-bottomed microtiter plates as single species, pairs, or triple species in root exudate. Plates are covered with sterile breathable sealing film and incubated at 20°C. Cell density was determined by spectrophotometric measurement at 600 nm at the final time point (BioTek Synergy HT). Planktonic cells were discarded, and wells were washed three times with water. Biofilms attached to the wells were stained with crystal violet 0.1%, washed three times with water, and dried. The stain was dissolved with 33% acetic acid, and its concentration was determined spectrophotometrically at 595 nm. Visualization and statistical analyses were performed with GraphPadPrism 7 software. Differences between groups were tested for statistical significance (Student's t-test). Significance levels were set to * P < 0.01.

---

**FIG S1.**
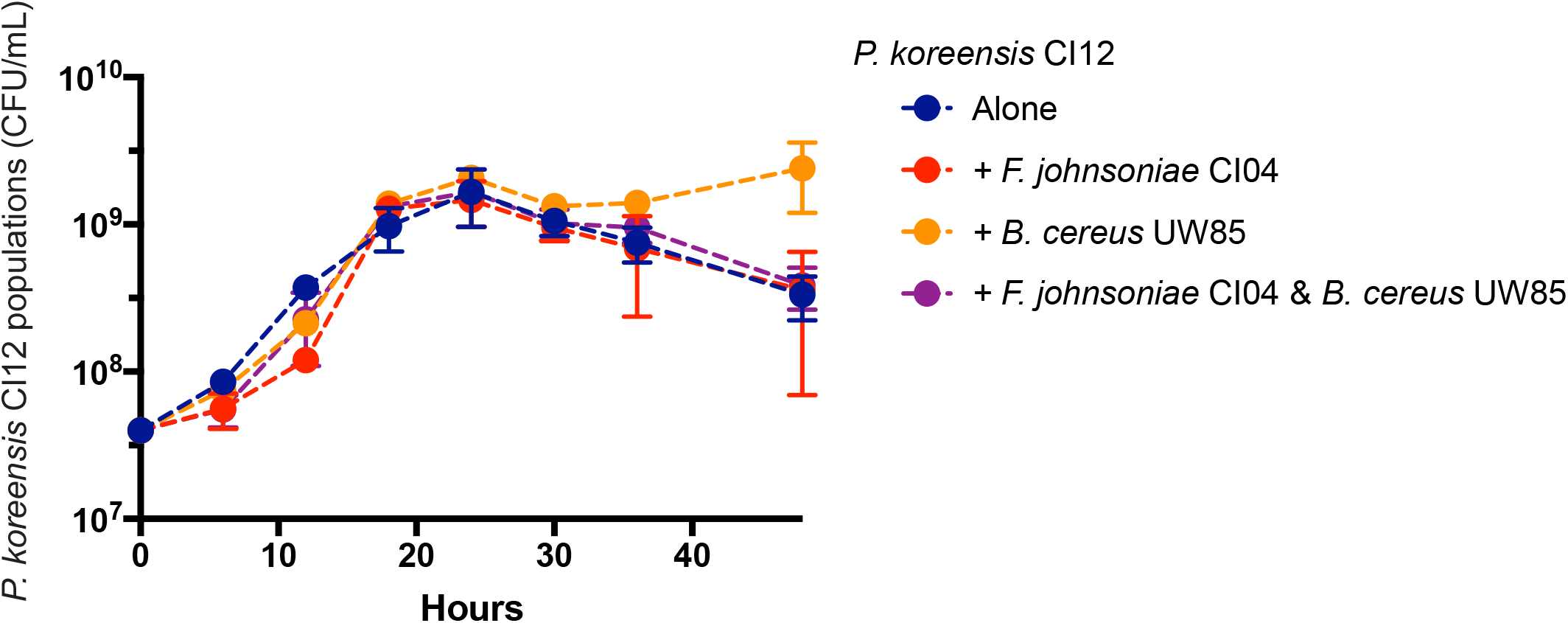
*P. koreensis* CI12 population dynamics. *P. koreensis* CI12 grown alone, in co-culture with *F. johnsoniae* CI04 or *B. cereus* UW85, and in triple culture with *F. johnsoniae* CI04 and *B. cereus* UW85.

**FIG S2.**
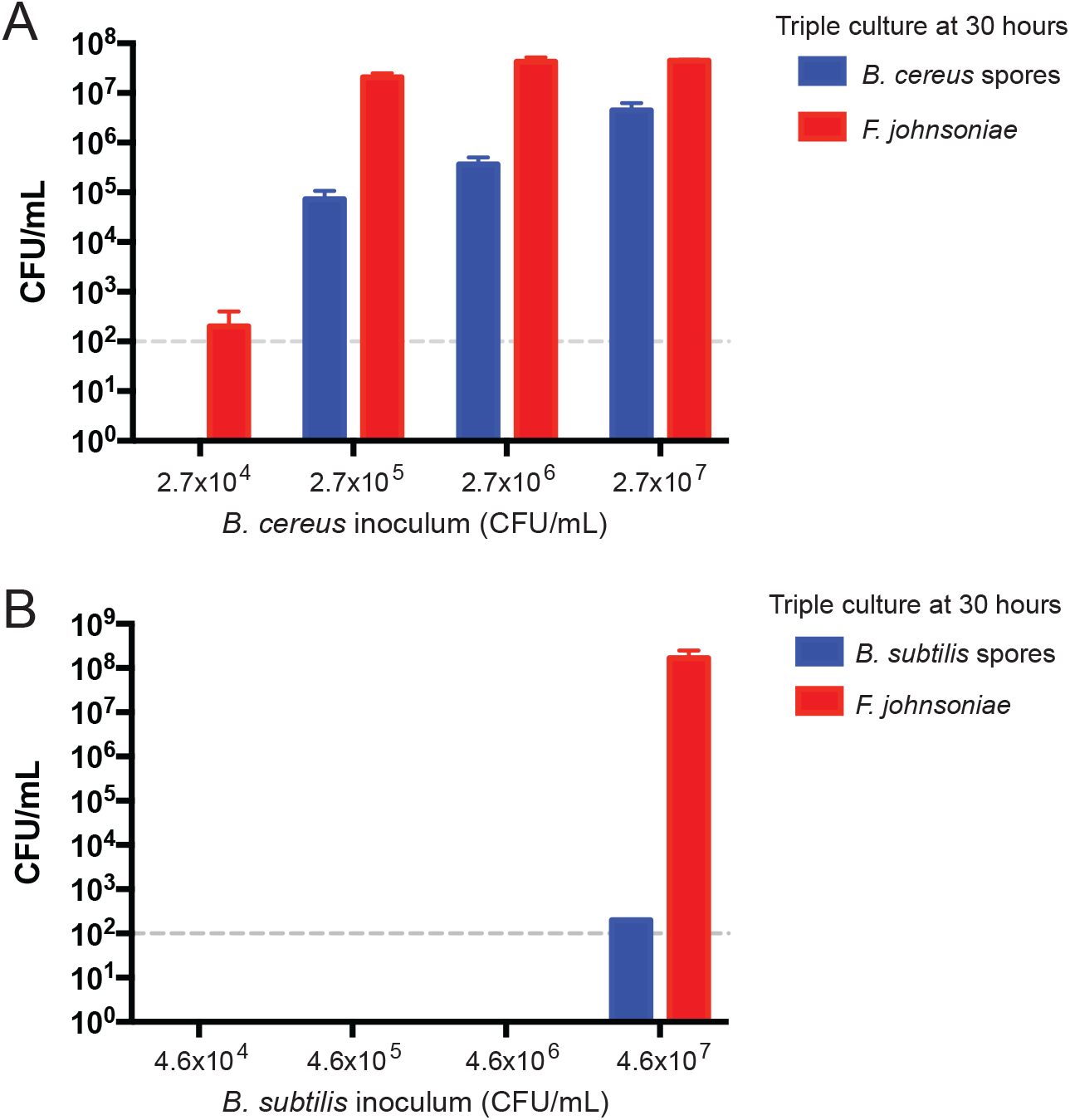
Correlation between *Bacillus spp.* spore density and *F. johnsoniae* protection in triple culture. Triple culture of *F. johnsoniae* CI04 and *P. koreensis* CI12 with either *B. cereus* UW85 (A) or *B. subtilis* NCIB3160 (B). The cell density of the *Bacillus* inoculum is on the x-axis. *Bacillus* spore densities and *F. johnsoniae* CI04 cell densities in the triple culture at 30 h are on the y-axis. Gray dotted line, limit of detection.

**FIG S3.**
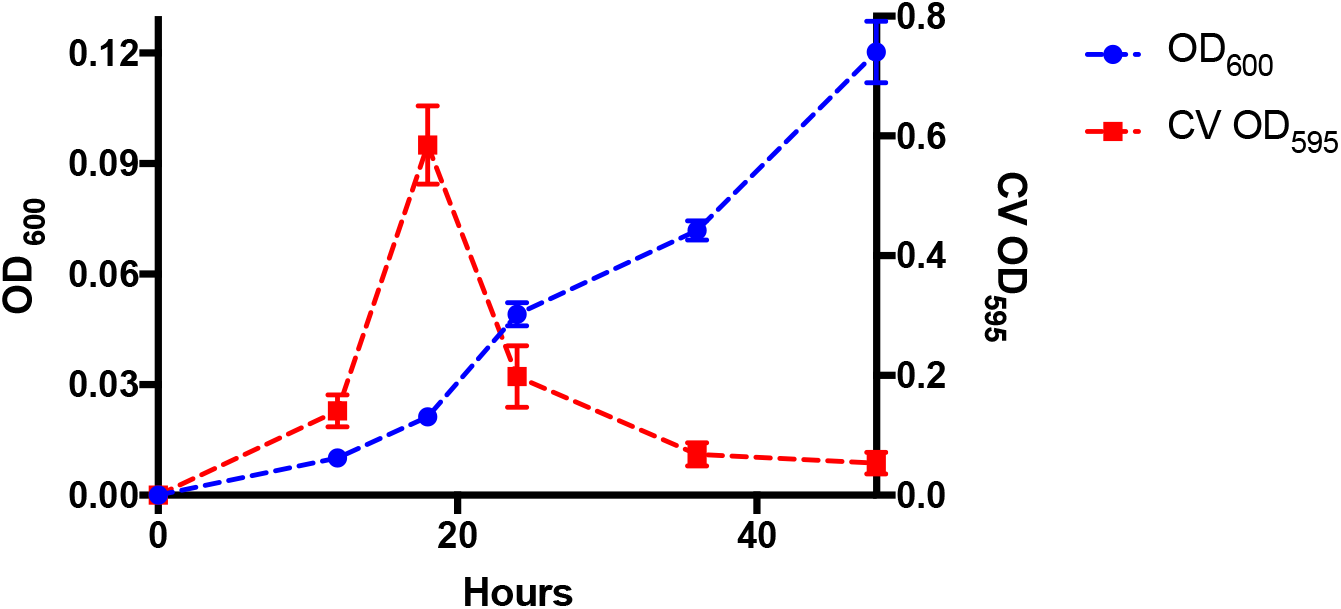
*P. koreensis* CI12 biofilm quantification and growth of the planktonic population. *P.* ! *koreensis* CI12 growth is on the left y-axis and crystal violet quantification of biofilm formation for *P. koreensis* CI12 is on the right y-axis.

**TABLE S1.** Area of *B. cereus* colonies is larger in the presence of either *F. johnsoniae* or *P. koreensis*.

|  | Area *B. cereus* colony (cm<sup>2</sup>) |  |
| --- | --- | --- |
|  | Day 2 | Day 5 |
| *B. cereus* alone | 1.3 +- 0.06 | 6.3 +- 0.42 |
| *B. cereus* with *F. johnsoniae* | 3.0 +- 0.24 | 24.3 +- 0.55 |
| *B. cereus* with *P. koreensis* | 1.8 +- 0.2 | 23.8 +- 2.66 |

**TABLE S2.**
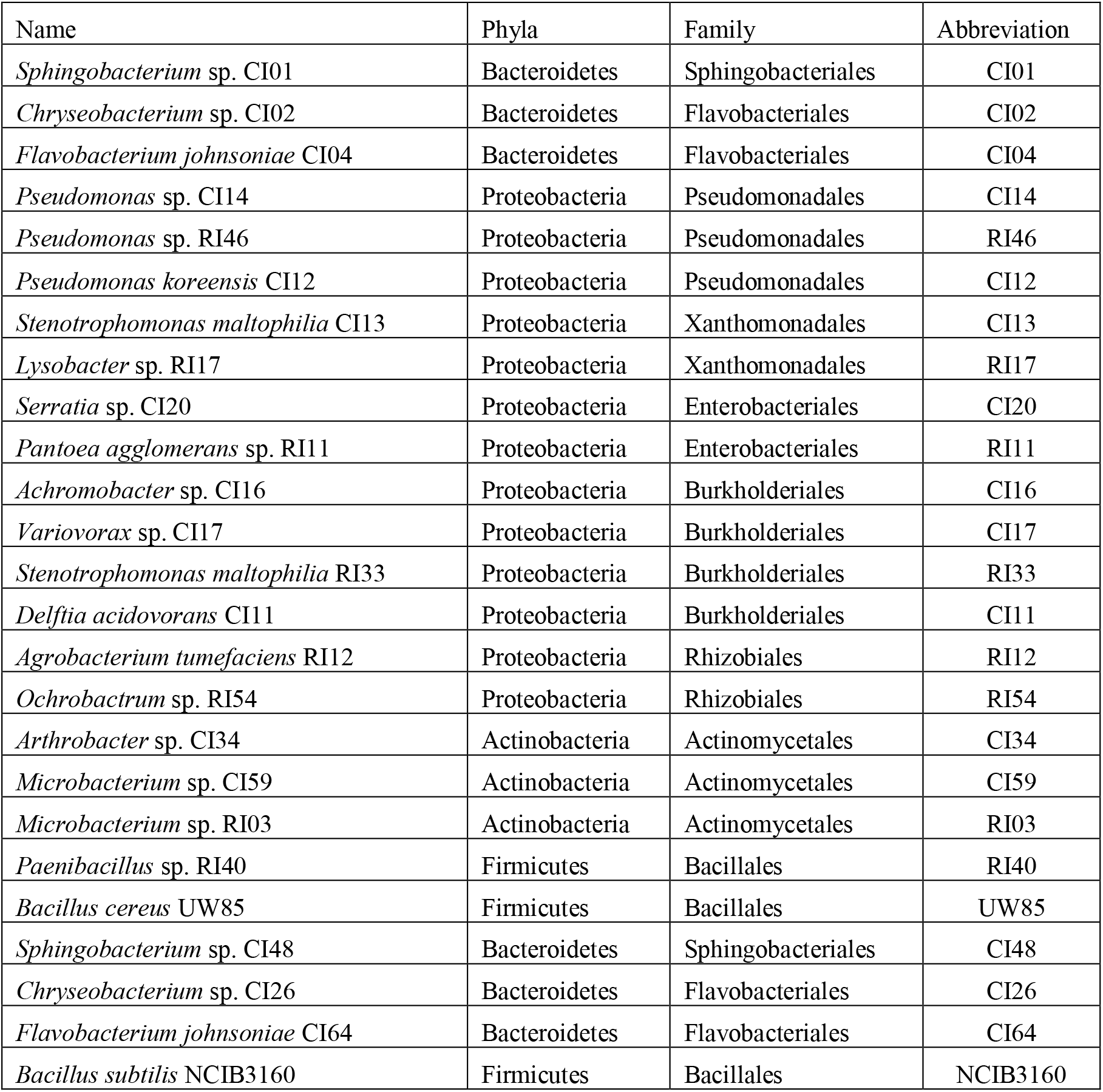
Strains used in this study.

**TABLE S3.**
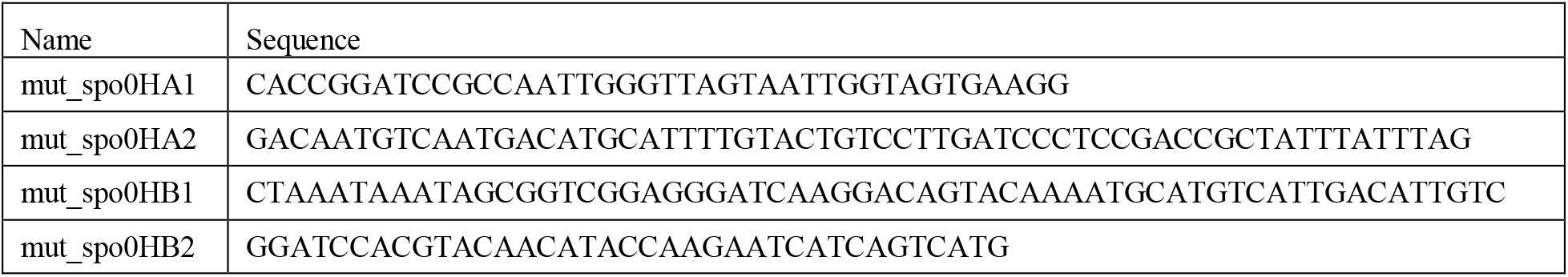
Primers used in this study.

